# Single-cell analysis of sterol-induced Ca^2+^ signaling in human astrocytes by dynamic mode decomposition

**DOI:** 10.64898/2026.02.09.704834

**Authors:** Miklas P. W. Larsen, Line Lauritsen, Rasmus Jensen, Ralf Zimmermann, Daniel Wüstner

## Abstract

Ca^2+^ signaling in astrocytes is a central mechanism of intercellular communication in the brain and plays a key role in regulating neuronal excitability, synaptic plasticity, and energy metabolism. Disruption of astrocytic Ca^2+^ dynamics is a characteristic of neurodegenerative diseases, as are deviations in cholesterol trafficking and metabolism, which are essential for maintaining membrane structure and function. Although recent studies have begun to explore links between Ca^2+^ signaling and sterol homeostasis in astrocytes, unbiased analytical workflows and mechanistic insight into how cholesterol and related sterols regulate astrocytic Ca^2+^ dynamics remain limited.

Here, we apply dynamic mode decomposition to dissect and classify Ca^2+^ signals obtained from time-lapse imaging of human astrocytes. Using both synthetic and experimental datasets, we show that delay-embedded dynamic mode decomposition combined with clustering separates heterogeneous Ca^2+^ activity into distinct dynamical states. This analysis reveals that increasing cholesterol levels shift astrocytes toward more active oscillatory states, whereas acute cholesterol depletion suppresses Ca^2+^ activity. In addition, pretreatment with the oxysterols 24-, 25-, and 27-hydroxycholesterol impaired cholesterol-induced Ca^2+^ oscillations.

Together, this work presents a general computational framework for decomposing and analyzing complex spatiotemporal Ca^2+^ signals, with broad applicability to quantitative imaging in cell biology.

## Introduction

Astrocytes are an abundant glial cell type in the Central Nervous System (CNS), and they play essential roles in brain function and intercellular communication (1, 2). Although they are electrically non-excitable, astrocytes display a form of cellular excitability through dynamic fluctuations in intracellular calcium ion (Ca^2+^) levels (3). These Ca^2+^ elevations activate downstream signaling pathways that allow astrocytes to modulate neuronal activity, vascular tone, and the behavior of neighboring glia cells (4–6). Through Ca^2+^-mediated signaling, astrocytes maintain brain homeostasis, and disruptions in these pathways are increasingly linked to neurological and neurodegenerative diseases including Alzheimer’s disease, Parkinson’s disease, and multiple sclerosis (3, 7–9).

Astrocytic Ca^2+^ signaling depends on membrane properties and ion channel function, both strongly influenced by lipid composition, particularly cholesterol, a key structural and regulatory component that modulates ion channel and receptor activity (10, 11). In the CNS, astrocytes are the principal source and distributor of cholesterol, and disturbances in astrocytic cholesterol metabolism are among the earliest features of neurodegenerative disorders (12, 13). Although cholesterol and its oxidized derivatives (oxysterols) are known to alter Ca^2+^ entry and membrane excitability in other cell types, their specific effects on astrocytic Ca^2+^ dynamics remain largely unexplored (14–16).

A defining characteristic of astrocytic Ca^2+^ signaling is its spatiotemporal heterogeneity at the single-cell level (17, 18). Individual astrocytes can exhibit transient single spikes, sustained Ca^2+^ elevations, low-frequency oscillations, or repetitive Ca^2+^ bursts, and these signals may propagate as intercellular Ca^2+^ waves across astrocytic networks (19–21). Even in uniform environments, astrocytes rarely behave the same way. One cell may generate rapid Ca^2+^ spikes, while the cell next to it shows only slow oscillations or none at all. This diversity stems from subtle differences in Ca^2+^-handling mechanisms as well as communication through gap junctions and extracellular messengers (20, 22, 23). Heterogeneous Ca^2+^ dynamics provide astrocytes with a versatile signaling range, but they also make quantitative analysis difficult at both the single-cell and population levels.

Understanding this heterogeneity requires large-scale singlecell Ca^2+^ imaging capable of tracking hundreds of astrocytes simultaneously. However, most conventional Ca^2+^ imaging studies are constrained by small fields of view and low-throughput acquisition, often the observation of only tens of cells at a time. These limitations obscure collective behaviors and mask the full distribution of single-cell responses within astrocyte populations. Moreover, standard analytical pipelines often rely on threshold-based peak detection or summary Δ*F/F*_0_ metrics, which discard the temporal structure, such as oscillation frequency, damping, or synchronization that distinguishes different Ca^2+^ signaling behaviors. Thus, there is a critical need for computational approaches that can decompose complex Ca^2+^ signals in an unbiased manner and classify distinct dynamical states across large astrocyte populations.

To overcome these limitations, we developed a large-scale single-cell Ca^2+^ imaging and computational analysis framework that enables quantitative characterization of astrocytic Ca^2+^ heterogeneity. Our pipeline employs widefield fluorescence microscopy with the highly sensitive Ca^2+^ indicator Cal-520 AM, providing high-throughput recordings from hundreds of cells per field with excellent signal-to-noise while minimizing dye-induced buffering effects. The superior brightness and retention of Cal-520, compared to conventional indicators such as Fluo-4, facilitate the reliable detection of subtle Ca^2+^ transients across extensive cell populations (24). After deep-learning-based segmentation of individual astrocytes from time-lapse recordings, we apply dynamic mode decomposition (DMD) to analyze Ca^2+^ signals. DMD is a data-driven matrix decomposition technique that provides a linear approximation of complex spatiotemporal dynamics (25). Originally developed in fluid dynamics, DMD has been successfully adapted for biomedical data analysis, including magnetic resonance imaging (MRI), positron emission tomography (PET), electroencephalogram (EEG) recordings, and live-cell fluorescence microscopy (26–37). Building upon these developments, we use a DMD variant to decompose and classify single-cell Ca^2+^ oscillations in human astrocytes subjected to controlled cholesterol loading, cholesterol depletion, and oxysterol treatment. This allows us to cluster cells into distinct groups with oscillation patterns for further analysis. Our high-throughput singlecell approach captures both common response patterns and rare, spatially localized events within astrocytic networks, enabling quantitative analysis of population heterogeneity.

Together, we present a novel imaging and computational framework in astrocytes that enables quantitative characterization of heterogeneous Ca^2+^ signaling dynamics and uncovers sterol-dependent remodeling of distinct Ca^2+^ activity states.

## Materials and Methods

### Cell Culture

Normal human astrocytes immortalized with SV40 large T antigen (P10251-IM, Innoprot) were maintained in T25 culture flasks (Nunc EasYFlask, 156367, Thermo Fisher Scientific) at 37 °C in a humidified incubator with 5% CO_2_. Cells were cultured in specialized astrocyte medium (Astrocyte Medium, AM; P60101, Innoprot), which was replaced every 2-3 days. Cell growth and contamination were monitored daily by light microscopy. Once the cell culture reached 90% confluency, cells were split by rinsing once with 1× phosphate-buffered saline (PBS; 70011044, Thermo Fisher Scientific) and incubating in 1× Trypsin-EDTA (T47174, Sigma-Aldrich) for 5 min at 37 °C. Detached cells were resuspended in fresh AM and transferred to new flasks.

### Material and Reagents

Cal-520 AM (21130, AAT Bioquest) was dissolved in DMSO Hybri-Max (D2650, Sigma-Aldrich) at 5 mM and stored in aliquots at −20 °C. Imaging medium (M1-media) consisted of 20 mM HEPES, 150 mM NaCl, 5 mM KCl, 1 mM CaCl_2_*·*2H_2_O, 1 mM MgCl_2_*·*6H_2_O, and 5 mM glucose, adjusted to pH 7.3 with NaOH and prepared using Milli-Q water. PBS was prepared by diluting 10× PBS to 1× using Milli-Q water.

Methyl-*β*-cyclodextrin (MCD; 332615, Sigma-Aldrich) was dissolved in 0.1% (w/v) bovine serum albumin (BSA; A3059, Sigma-Aldrich) prepared in 1× PBS. To generate cholesterol-cyclodextrin complexes, cholesterol (C8667, Sigma-Aldrich) was combined with MCD at a molar ratio of 1:8. The mixture was sonicated for 20 minutes on ice to facilitate solubilization, followed by centrifugation at 20,000× *g* for 20 minutes at 4°C. The resulting supernatant, containing water-soluble cholesterol-cyclodextrin complexes, was collected and used for subsequent experiments. Dehydroergosterol (DHE; E2634, Sigma-Aldrich) was prepared as a water-soluble cyclodextrin complex following the same protocol used for cholesterol-cyclodextrin complexes. 24(S)-Hydroxycholesterol (SML1648, Sigma-Aldrich), 25-hydroxycholesterol (H1015, Sigma-Aldrich), and 27-hydroxycholesterol (700021, Avanti) were dissolved in absolute ethanol at 5 mM. Oxysterols were diluted and added directly to astrocytes at a final concentration of 10 µM.

### Calcium Imaging

For calcium imaging, astrocytes were seeded into µ-Slide 8-well chambered coverslips (80826, Ibidi) and cultured to ∼70% confluency. Cells were washed once with 1× PBS and incubated with 0.5 µM Cal-520 AM in M1-media for 40 minutes at 37 °C in 5% CO_2_. After incubation, cells were washed three times with 1× PBS and replenished with fresh M1-media. Live imaging was performed using a Nikon Ti2 Eclipse inverted widefield microscope (Nikon Corporation, Tokyo, Japan) equipped with a CoolLED pE-300 Illumination System and an Andor Zyla 5.5 MP sCMOS camera (Oxford Instruments, Belfast, UK). Temperature was maintained at 37 °C using an H301-NIKON-TI-S-ER stage-top incubator in conjunction with an OKOLab UNO controller (OKOLab, Pozzuoli, Italy). Cells were allowed to equilibrate on the microscope stage for 15 minutes prior to image acquisition. Images were acquired using a 20×/0.75 NA air objective lens and a FITC filter cube with 488 nm excitation. Unless otherwise specified, acquisition was performed using 2% excitation intensity, 100 ms exposure time, and a capture interval of 1 second for 20 minutes.

### Image segmentation and extraction of time series

Maximum intensity projections were generated from each time-lapse image stack using FIJI (https://imagej.net/). Cell segmentation was performed using the deep-learning-based algorithm Cellpose (38) with a pre-trained model optimized for in vitro astrocytes based on cyto 2 model with settings of diameter = 100 pixels and flow threshold = 0.4. The resulting segmentation masks were used to extract mean fluorescence intensity values for each individual cell across all time points. To quantify calcium activity, fluorescence time series from each cell were normalized using a rolling baseline normalization method (ΔF/F_0_). For each time point, F_0_ was defined as the minimum fluorescence intensity within a window of 50 frames before and 50 frames after the time point of interest. The normalized signal was then calculated as seen in Eq. (1), where F represents the raw fluorescence intensity. This approach allowed for detection of calcium dynamics while minimizing the effect of photobleaching and accounting for heterogeneous intensities. Peaks were identified using the `find_peaks` function from the `scipy.signal` library (39). Peaks were detected on the ΔF/F_0_ signal using a minimum height of 0.2 and a prominence threshold of 0.1. For each detected peak, the time of occurrence, amplitude, and full width at half maximum (FWHM) were extracted and used for descriptive analysis of Ca^2+^ activity across conditions.

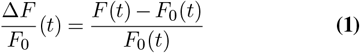

### Time series analysis

#### Dynamic mode decomposition

The snapshot matrix of cellular calcium traces, *X*, was concatenated with time-shifted versions of each data set to form a Hankel, *H*, matrix as described (37):

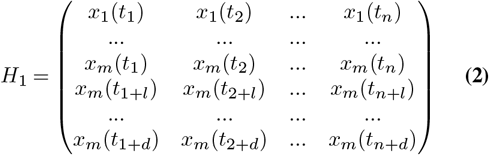

Here, instances of time, at which the fluorescence of the calcium was measured are *t*_*k*_, while the corresponding calcium signal for cell *i* at this time point is *x*_*i*_(*t*_*k*_). The index *l* is the time delay (chosen as *l* = 1) and *d* is the embedding dimension, i.e., the total number of chosen time shifts. DMD carried out on this Hankel matrix seeks to find a linear operator, *A*, describing the progression from state *k* to state *k*+1 according to:

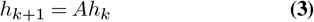

To obtain *A* one minimizes the Frobenius norm of

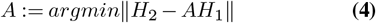

This requires finding the Moore-Penrose pseudoinverse of *H*_1_, for which one uses a singular value decomposition of the (delay-embedded) snapshot matrix *H*_1_ into orthogonal matrices *U* and *V\** and diagonal matrix Σ holding the singular values of *H*_1_. The Hankel matrix *H*_2_ is the time-shifted version of matrix *H*_1_, i.e. one step forward in time. DMD is intended to give a low-rank approximation of the matrix *A*, up to rank *r*<(*m*-1), which is obtained by projecting it onto the left singular vectors (i.e. the column vectors of *U*):

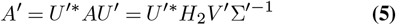

Spectral decomposition of the rank-truncated system matrix *A’* gives the solution

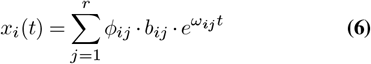

Here, *x*_*i*_(*t*) is the reconstructed image time series for the *i*th cell, *b*_*ij*_ the initial value constant, *ϕ*_*ij*_ the eigenfunction (DMD mode) and *ω*_*ij*_=ln(*λ*_*ij*_*/δt*) the corresponding eigenvalue of mode *j* of cell *i*. For practical reasons, we have carried out the DMD analysis for all cells in a data matrix at once, resulting in identical eigenvalues for all cells in a given mode, i.e. *ω*_*ij*_=*ω*_*j*_. Since the clustering is based on the (cell- and mode-specific) eigenfunctions, *ϕ*_*ij*_, this approach is appropriate for both, synthetic and experimental data. Comparable reconstruction quality was achieved, when performing the DMD analysis on each time series and its delay-embedded version separately on synthetic data (not shown, but see Fig S1).

#### Kernel PCA and cluster analysis

DMD amplitudes were calculated as the square root of the sum of the real and imaginary parts of the eigenfunctions (i.e. of the DMD modes). Classification of these amplitudes was achieved using k-Means clustering of the feature space obtained by kernel principal component analysis, as described (37). The optimal number of Principal Components were determined based on the Silhouette and Elbow criteria implemented in the Python library scikit-learn (40).

#### Analysis of synchronization

From the cell cluster matrices *X*_*n*_ ∈ ℝ^*N* ×*T*^, for clusters *n* = 1 … 3 identified by DMD and kernel-based time series classification, temporal correlation matrices were computed. All traces were high-pass filtered using a second-order Butterworth filter with a cutoff frequency of 0.005 Hz, applied with zero-phase forward–backward filtering to remove slow baseline fluctuations and quiescent activity while preserving temporal phase relationships. For each cluster, activity was z-scored across cells at each time point and the temporal correlation matrix was calculated as

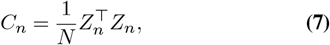

where *Z*_*n*_ denotes the normalized activity matrix.

To quantify phase synchronization, the instantaneous phase *ψ*_*i*_(*t*), amplitude, and frequency of each cell were extracted from the analytic signal obtained via the Hilbert transform of each time series *x*_*i*_(*t*),

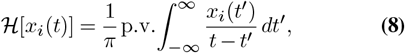

where p.v. denotes the Cauchy principal value. The instantaneous phase was obtained as the argument of the analytic signal.

From the phases of the *N* cells within each cluster, the Kuramoto order parameter was computed as (41, 42)

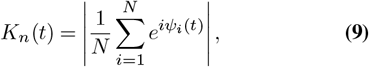

which quantifies the degree of instantaneous phase synchronization.

## Results

### DMD and time series classification of simulated Ca^2+^ dynamics

Synthetic data are often used to validate the performance of a machine learning algorithm, and we used a simple and well-perceived model of Ca^2+^ oscillations in astrocytes to assess the performance of our approach. This model is based on non-linear ordinary differential equations (ODEs) and models the core elements being required to generate Ca^2+^ oscillations in astrocytes (43, 44). It includes (1) inositol-1,4,5-trisphosphate receptors (IP_3_R) in the endoplasmic reticulum (ER), mediating IP_3_-dependent Ca^2+^ release into the cytosol, (2) the SERCA pump, which returns Ca^2+^ from the cytosol back into the ER in an ATP-dependent manner, and (3) phospholipase C (PLC)–mediated production of IP_3_ and diacylglycerol (DAG) from phosphatidylinositol-4,5-bisphosphate (PIP_2_), initiated by activation of a Gprotein–coupled receptor (GPCR) at the plasma membrane (Fig. 1A). Given that previous studies implicated the activity of the SERCA pump in cholesterol-induced Ca^2+^ oscillations in astrocytes (15), we simulated that model by numerical integration of the underlying ODE system for varying SERCA activity (Fig. 1B). To generate realistic conditions, noise was added to each time course (see Materials and methods for details). Clearly, varying SERCA activity affects the nature of the oscillations; for low activity, simulated with *v*_*m*2_, the rate for SERCA being equal to 5 µM /s, only damped oscillations are found, which lead to a stable node in the phase plot (green line in Fig. 1B-D). For higher SERCA activity sustained oscillations are found, which correspond to a limit cycle in the phase plot (orange and blue lines in Fig. 1B-D). In this range of SERCA activity of *v*_*m*2_ = 10-15 µM/s, the extent of Ca^2+^ pumping back into the ER controls the phase and frequency of Ca^2+^ spiking, in accordance with the original study ((45). To generate single-cell data with stochastically varying Ca^2+^ activity we generated 100 random samples from a Gaussian distribution of SERCA activity (Fig. 1E and Fig. 2A). Clearly, stochastic variation of SERCA activity around three distinct mean values generates characteristic oscillation patterns, which serve as input for the DMD analysis.

**Fig. 1:**
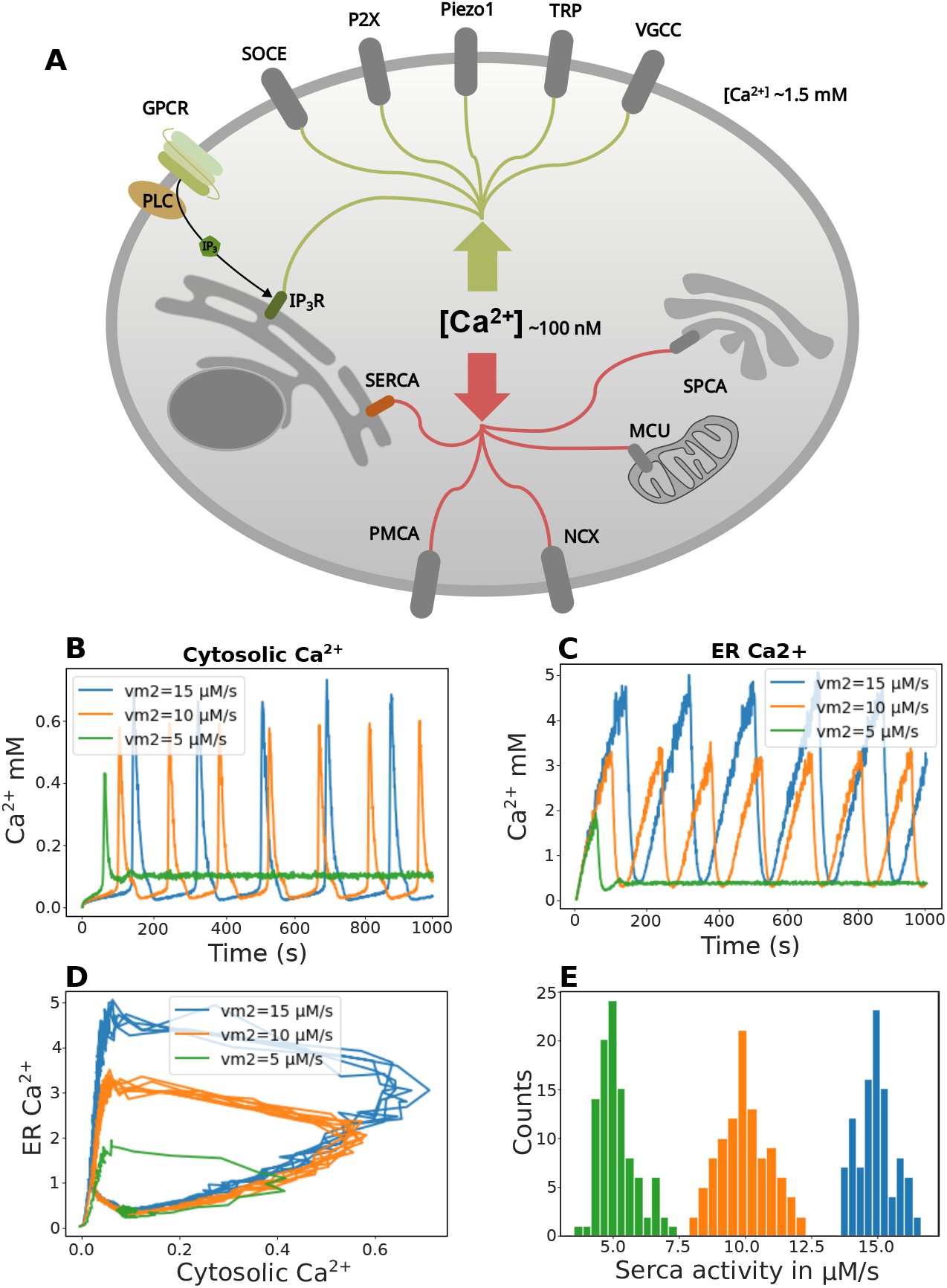
Simulation of calcium oscillations in astrocytes under varying SERCA pump activity. (A) Schematic representation of intracellular calcium handling components, illustrating selected proteins involved in astrocytic calcium signaling. The ordinary differential equation (ODE)–based framework incorporates cytosolic calcium influx pathways, inositol 1,4,5-trisphosphate receptor (IP_3_R)-mediated endoplasmic reticulum (ER) calcium release (Green), and SERCA-driven reuptake (Red), while other signaling modules are greyscale for simplicity. Components shown include G-protein-coupled receptors (GPCRs), phospholipase C (PLC), store-operated calcium entry (SOCE), purinergic P2X receptors (P2X), Piezo1 channels, transient receptor potential (TRP) channels, voltage-gated calcium channels (VGCCs), plasma membrane Ca^2+^-ATPase (PMCA), Na^+^/Ca^2+^ exchanger (NCX), mitochondrial calcium uniporter (MCU), and secretory pathway Ca^2+^-ATPase (SPCA). (B) Simulated cytosolic Ca^2+^ dynamics for three levels of SERCA activity (*v*_*m*2_ = 5, 10, and 15 *µ*M/s). Higher SERCA rates lead to more frequent and higher-amplitude oscillations. (C) Corresponding ER Ca^2+^ dynamics show complementary depletion and refilling behavior across the same SERCA conditions. (D) Phase-plane trajectories (ER Ca^2+^ vs. cytosolic Ca^2+^) illustrate distinct dynamical regimes that emerge as a function of SERCA strength. (E) Distribution of 100 sampled *v*_*m*2_ values drawn from three Gaussian distributions centered at 5 *µ*M/s (green), 10 *µ*M/s (orange), and 15 *µ*M/s (blue), used to generate the simulated variability in Ca^2+^ dynamics. Together, the simulations demonstrate how altering SERCA activity modulates the frequency, amplitude, and qualitative form of astrocytic calcium oscillations.

**Fig. 2:**
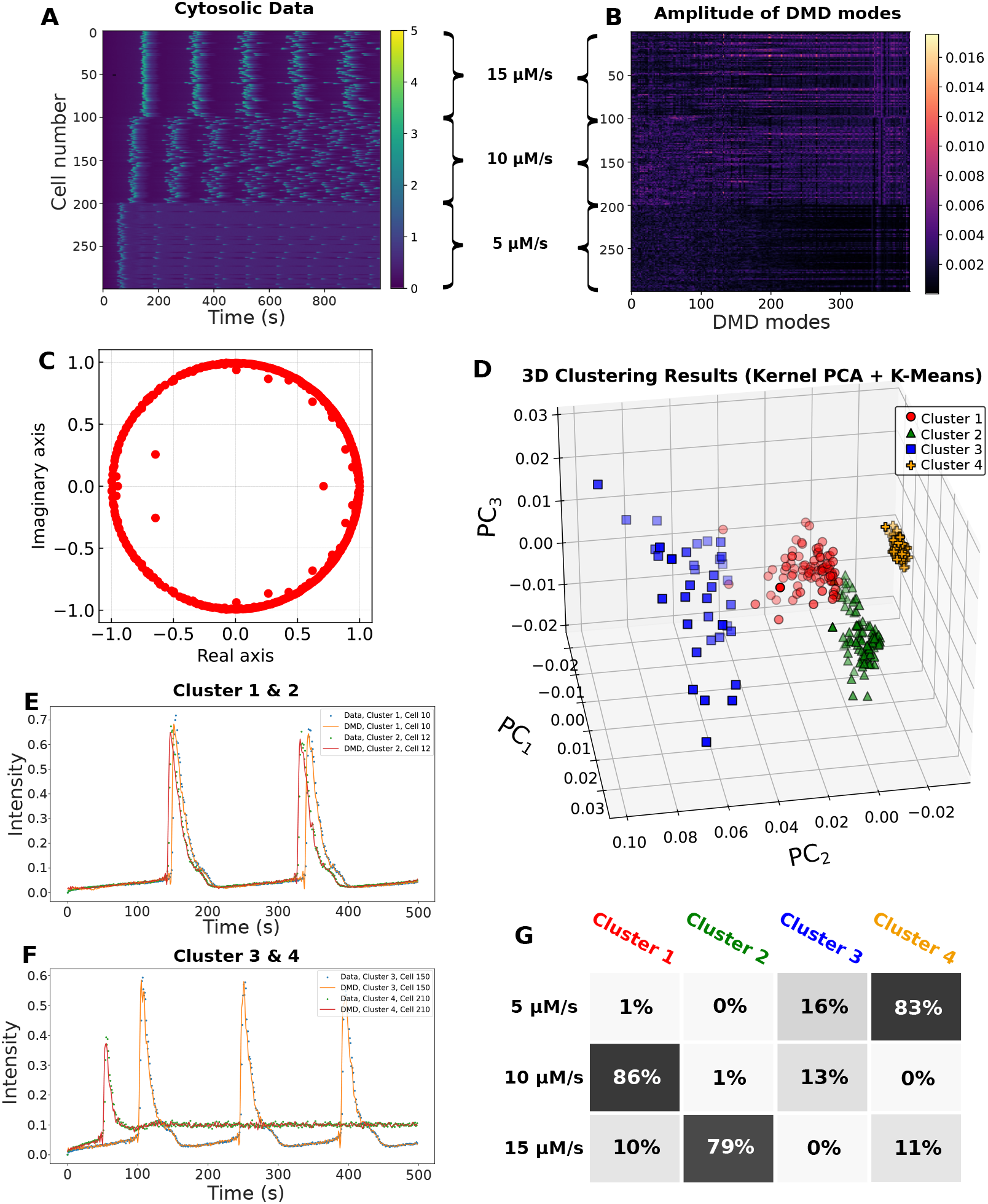
DMD reconstruction and classification of simulated single-cell Ca^2+^ dynamics. (A) Cytosolic Ca^2+^ traces generated by numerically integrating the astrocyte ODE model for three values of the SERCA rate parameter *v*_*m*2_, producing 100 simulated cells per condition. (B) Amplitudes of the DMD modes and (C) corresponding eigenvalues computed from a rank-400 decomposition with a delay dimension of 300. (D) Kernel PCA applied to the DMD mode amplitudes followed by *k*-means clustering separated the simulated cells into four dynamical clusters. Silhouette score = 0.64 (E–F) Example Ca^2+^ traces (dots) and their DMD reconstructions (lines) for cells representative of clusters 1–2 (E) and clusters 3–4 (F). (G) Cluster composition across experimental conditions indicating the distribution of cells in each cluster.

Classification of the highly non-linear Ca^2+^ dynamics at a single-cell level requires to derive a locally linear approximation of the dynamics, and we aimed for using DMD for this purpose. Direct application of DMD to the simulated singlecell time courses, however, was unsuccessful, as there is no linear subspace, which approximates the highly non-linear and oscillatory dynamics adequately (not shown, but see (37) for a similar problem). We therefore generated a Hankel matrix H of the simulated snapshot series, which contains time-shifted (delayed) versions of the data, thereby enriching the sample information about the past for each of the trajectories (see Materials and methods, Fig. S1 and (37) for details. The delay-embedded snapshot matrix can be seen as a function dictionary for DMD, enabling better linear approximation of the dynamics within this function space (46). We used a delay of d = 300 and a rank, r=400, truncation of the DMD matrix, A, approximating the Koopman operator in a linear subspace (Fig. 2B). The eigenvalue spectrum and eigenfunctions (DMD modes) contain the temporal and spatial information of the decomposition, respectively Fig. 2C and D). Using just the DMD modes, each cell is represented as a data point in a 400-dimensional space, and we use kernel-PCA for dimensionality reduction followed by clustering using a k-means algorithm, as recently described (37, 47). This procedure results in four clusters, which nicely dissect the simulated data, and efficiently classifies the trajectories according to their oscillation behavior reflecting varying SERCA activity Fig. 2E). Importantly, clustering into four groups was selected as optimal based on evaluation using the silhouette score and the elbow method (not shown)

### DMD-based time series classification of cholesterol-induced Ca^2+^ oscillations in astrocytes

Next, we used the same approach on experimental Ca^2+^ data, here immortalized human astrocytes were treated with methyl-betacyclodextrin (MCD) to remove cellular cholesterol or with cholesterol-cyclodextrin complexes to load cells with cholesterol. To obtain reliable single-cell Ca^2+^ traces, we first established a Ca^2+^ imaging pipeline based on the Cal-520 AM sensor, which was found to be the most sensitive Ca^2+^ indicator (Fig. 3). We also evaluated several imaging conditions, including temperature, light intensity, and exposure time, and found that recordings performed at 37 °C with low excitation power and short exposure times produced the most stable and biologically accurate Ca^2+^ signals (Fig S2).

**Fig. 3:**
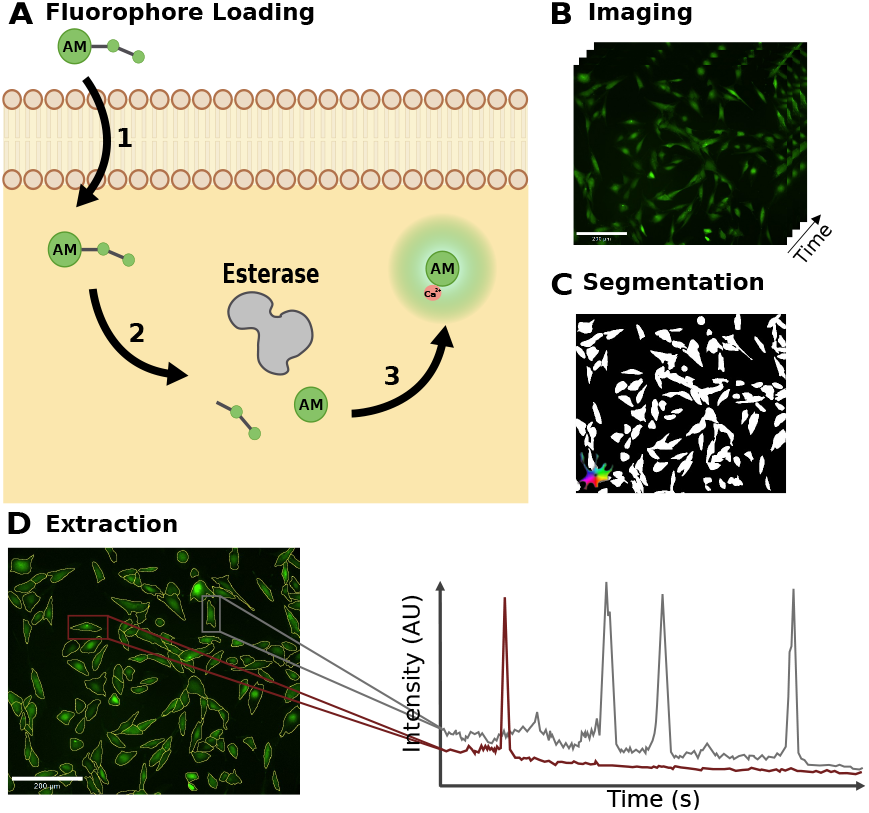
Schematic overview of experimental Ca^2+^ imaging utilizing fluorophore loading and extraction of single-cell Ca^2+^ traces. (A) Fluorophore loading: (1) Cells were incubated with acetoxymethyl ester Ca^2+^ indicators, which passively diffuse across the plasma membrane. (2) Intracellular esterases hydrolyze the AM groups, thereby trapping the fluorophore in the cytosol. (3) Upon binding to intracellular Ca^2+^, the indicators increase in fluorescence intensity, enabling visualization of Ca^2+^ dynamics. (B) Time-lapse fluorescence microscopy was used to record Ca^2+^ activity in vitro. (C) Image stacks were processed and segmented into individual cells using pretrained models in Cellpose. (D) Fluorescence intensity was extracted for each segmented region of interest over time to generate single-cell Ca^2+^ traces.

Cells were then imaged on a widefield fluorescence microscopy at 1 Hz for 20 minutes. After recording, the time-lapse stacks were segmented using a pre-trained Cellpose model, and fluorescence time series were extracted from each identified cell to generate the single-cell Ca^2+^ traces used for DMD analysis (Fig. 3B-D). All treatments were added at *t* = 300 s during live imaging.

Under control conditions, astrocytes exhibited spontaneous Ca^2+^ oscillations with occasional spikes. Acute cholesterol depletion using MCD markedly reduced this activity, whereas cholesterol loading increased both the amplitude and frequency of Ca^2+^ transients, most prominently at 100 and 250 µM cholesterol–cyclodextrin (Fig. 4A and Fig. S3).

**Fig. 4:**
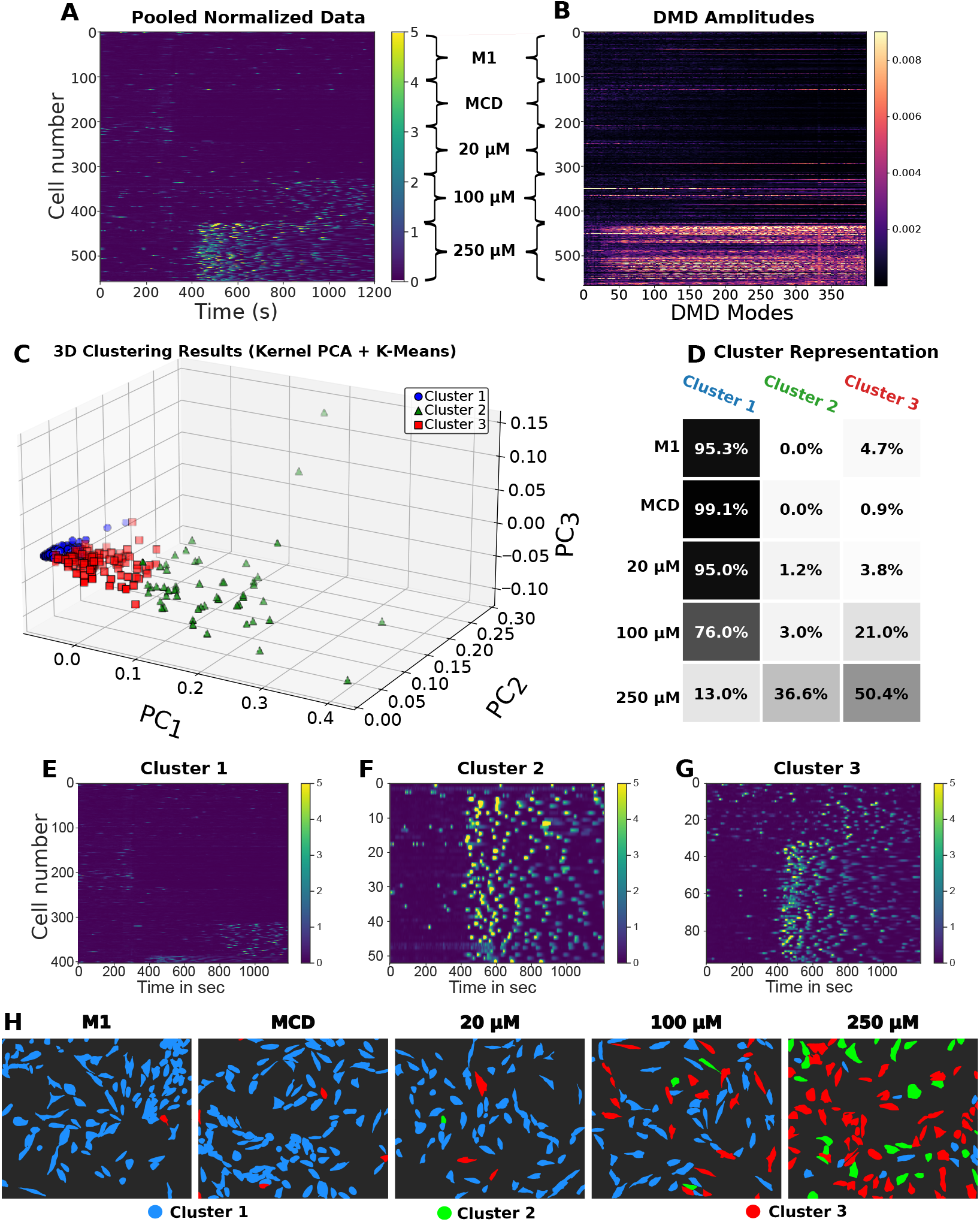
Classification of cholesterol-dependent single-cell Ca^2+^ dynamics using DMD-TDE and kernel-PCA. (A) Pooled and normalized cytosolic Ca^2+^ traces from all experimental conditions. Astrocytes were loaded with Cal-520 and imaged at 1 Hz; at *t* = 300 s, either control medium (M1), methyl-beta-cyclodextrin (MCD), or cholesterol–cyclodextrin complexes (20, 100, or 250 µM cholesterol) were added. Traces from all cells across all conditions were concatenated into a single cell-time matrix. (B) Amplitudes of the delay-embedded DMD modes computed from the full dataset (delay embedding: 400 steps; truncation rank: 400). (C) Kernel-PCA followed by *k*-means clustering identified three distinct dynamical clusters of Ca^2+^ activity. Silhouette score = 0.708 (D) Cluster representation across conditions, showing strong enrichment of cluster 1 in control and cholesterol-depleted cells, and a shift toward clusters 2 and 3 with increasing cholesterol concentration. (E–G) Ca^2+^ heatmaps (identical intensity scaling) for the three clusters, illustrating that cluster 1 exhibits low activity, cluster 2 displays frequent high-amplitude spikes, and cluster 3 shows dense, moderate-amplitude oscillations. (H) Spatial maps of classified cells for each experimental condition. Cells are color-coded according to their assigned cluster (blue: cluster 1; green: cluster 2; red: cluster 3), and cell outlines are derived from CellPose segmentation (see Materials and Methods).

To verify that sterols were rapidly delivered to cells under the same experimental conditions, we monitored sterol incorporation using a cyclodextrin–dehydroergosterol (DHE) complex. DHE fluorescence became detectable at the plasma membrane and in intracellular compartments within approximately 2 minutes of addition, preceding the onset of Ca^2+^ responses (Fig. S4). These observations confirm that the observed changes in Ca^2+^ dynamics occur after sterol delivery to cells.

DMD with time-delay (DMD-TDE) embedding revealed clear differences in the DMD modes, particularly at the highest cholesterol-cyclodextrin concentration (250 µM cholesterol) (Fig. 4B). Kernel PCA followed by *k*-means clustering separated the single-cell Ca^2+^ trajectories into three distinct dynamical clusters (Fig. 4C). Cluster 1 (blue symbols) consisted predominantly of control (M1), cholesterol-depleted (MCD), and low cholesterol-cyclodextrin (20 µM) treated cells, whereas clusters 2 and 3 (green and red symbols) contained most cells exposed to elevated cholesterol concentrations (100-250 µM). This distribution is reflected in both the cluster representation matrix and the traditional peak-based analysis (Fig. 4D; Fig. S5).

Inspection of the Ca^2+^ activity patterns assigned to each cluster reveals that the clusters exhibit markedly different dynamical signatures (Fig. 4E-G). Cluster 1 cells show weak or absent activity with mostly flat traces, consistent with quiescent Ca^2+^ behavior. Cluster 2 cells display sporadic high-amplitude spiking events, whereas cluster 3 cells show frequent but lower-amplitude oscillations. These differences are also visible in the representative spatial maps (Fig. 4H), which show that increasing cholesterol levels lead to a clear redistribution of cells among the identified clusters. This shows that the DMD-TDE approach captures distinct Ca^2+^ activity patterns that emerge as astrocytes respond to changes in cholesterol.

### Cluster-specific coordination and synchronization of astrocytic Ca^2+^ dynamics

Apart from adapting the frequency and amplitude of their Ca^2+^ spikes, astrocytes can also respond to changing conditions by synchronizing or desynchronizing their Ca^2+^ dynamics. In vivo and in 2D cell culture, coordinated Ca^2+^ activity has been observed among groups of neighboring astrocytes, often mediated by metabotropic glutamate receptor activation (48, 49). Such collective Ca^2+^ signaling is thought to contribute to astrocyte–astrocyte communication and may modulate neuronal network states, including slow-wave activity (50–52).

To assess the degree of synchronization within each cluster identified by our clustering pipeline, we applied two complementary analyses. First, we computed temporal correlation matrices for each cluster after applying a high-pass filter to suppress quiescent activity that would otherwise dominate synchronization measures (Fig. 5A–C). These matrices reveal stronger off-diagonal structure in clusters 2 and 3 compared to cluster 1, indicating the presence of recurrent or coordinated Ca^2+^ events. Moreover, the distinct correlation patterns observed in clusters 2 and 3 suggest qualitatively different modes of coordinated activity.

**Fig. 5:**
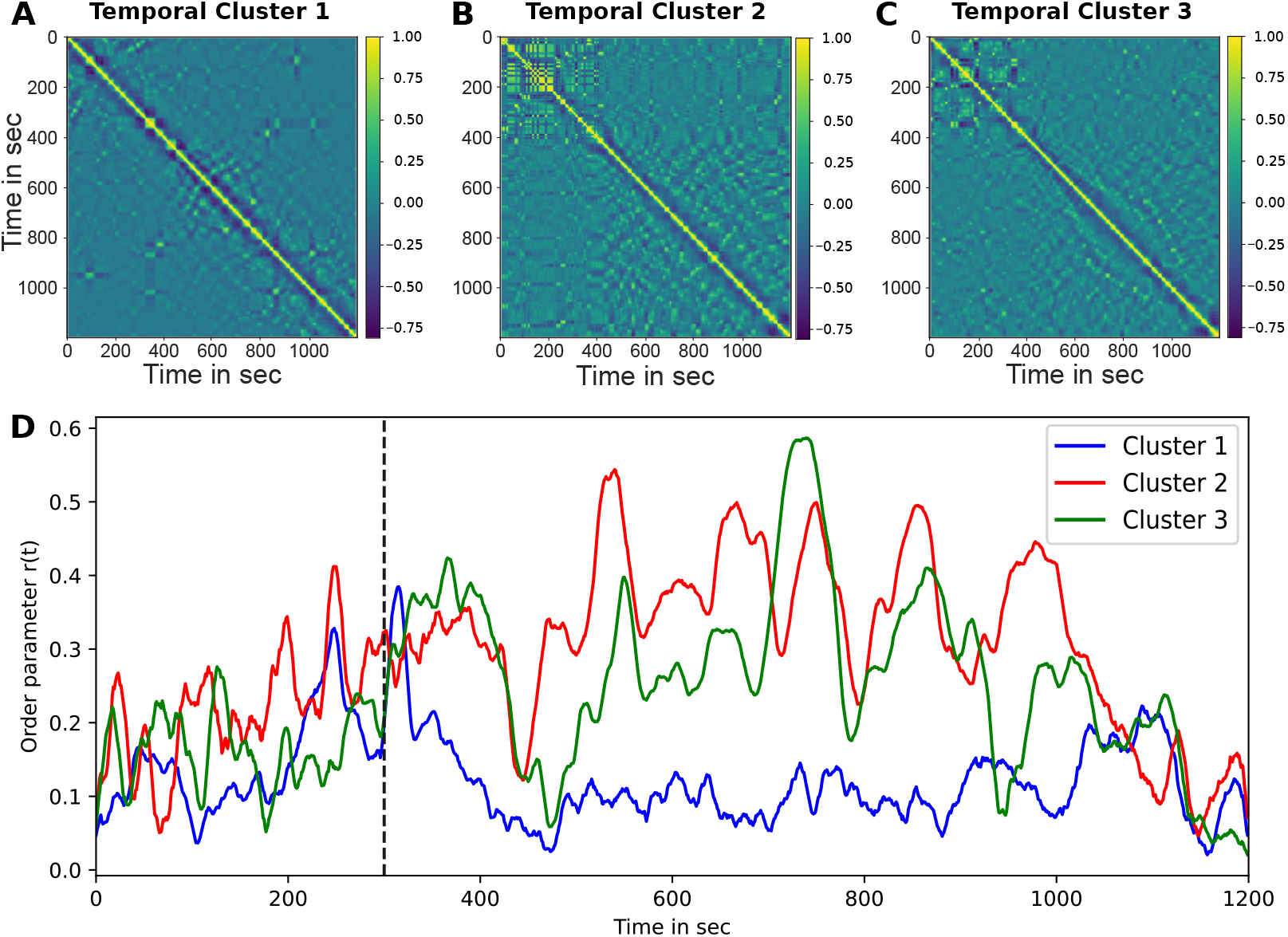
Temporal correlation structure and synchronization in the three astrocyte clusters. (A-C) Temporal correlation matrices for clusters 1-3, computed from the Ca^2+^ activity time series within each cluster. Clusters 2 and 3 show pronounced off-diagonal structure, indicating recurrent and coordinated activity patterns over time, whereas cluster 1 exhibits minimal off-diagonal correlations, consistent with weak temporal coupling. (D) Kuramoto order parameter *r*(*t*) for each cluster, quantifying instantaneous phase synchrony. The dashed vertical line marks the addition of cholesterol-cyclodextrin complexes at 300 s.

To further quantify synchronization of Ca^2+^ oscillations, we extracted the instantaneous phase, amplitude, and frequency of each cell using the Hilbert transform (see Materials and Methods) (42). From the phase information, we computed the Kuramoto order parameter, which approaches one for perfectly synchronized populations and zero for completely incoherent spiking behavior. Following cholesterol addition, clusters 2 and 3 exhibited pronounced, transient increases in the order parameter, reflecting enhanced collective signaling (Fig. 5D). Notably, both clusters showed discrete synchronization events, with cluster 2 displaying more frequent and sustained peaks in the order parameter.

In contrast, cluster 1 comprising predominantly untreated and cholesterol-depleted cells showed low and weakly structured synchronization after treatment, with the order parameter fluctuating around values of approximately 0.1 (Fig. 5D, blue trace).

### Oxysterols reshape astrocytic Ca^2+^ responses to rapid cholesterol loading

Oxysterols are potent regulators of membrane organization, ion channel activity, and lipid-sensitive signaling pathways, and their levels increase in several neurological diseases (53, 54). Because oxysterols can influence Ca^2+^ entry and ER Ca^2+^ handling in other cell types, we asked whether prolonged exposure to hydroxycholesterols alters the astrocytic Ca^2+^ response to acute cholesterol loading (14, 55). To address this, astrocytes were pretreated for 24 hours with 24-HC, 25-HC, or 27-HC prior to the addition of 250 µM cholesterol-cyclodextrin.

In control cells, acute cholesterol loading induced a pronounced and coordinated Ca^2+^ elevation beginning approximately 80 s after cholesterol addition, followed by sustained oscillatory activity (Fig. S6 and Fig. 6A). In contrast, pretreatment with any of the three oxysterols strongly attenuated this response. The large, collective Ca^2+^ spikes observed in control cells were largely absent, and subsequent oscillatory activity was markedly reduced or completely suppressed.

**Fig. 6:**
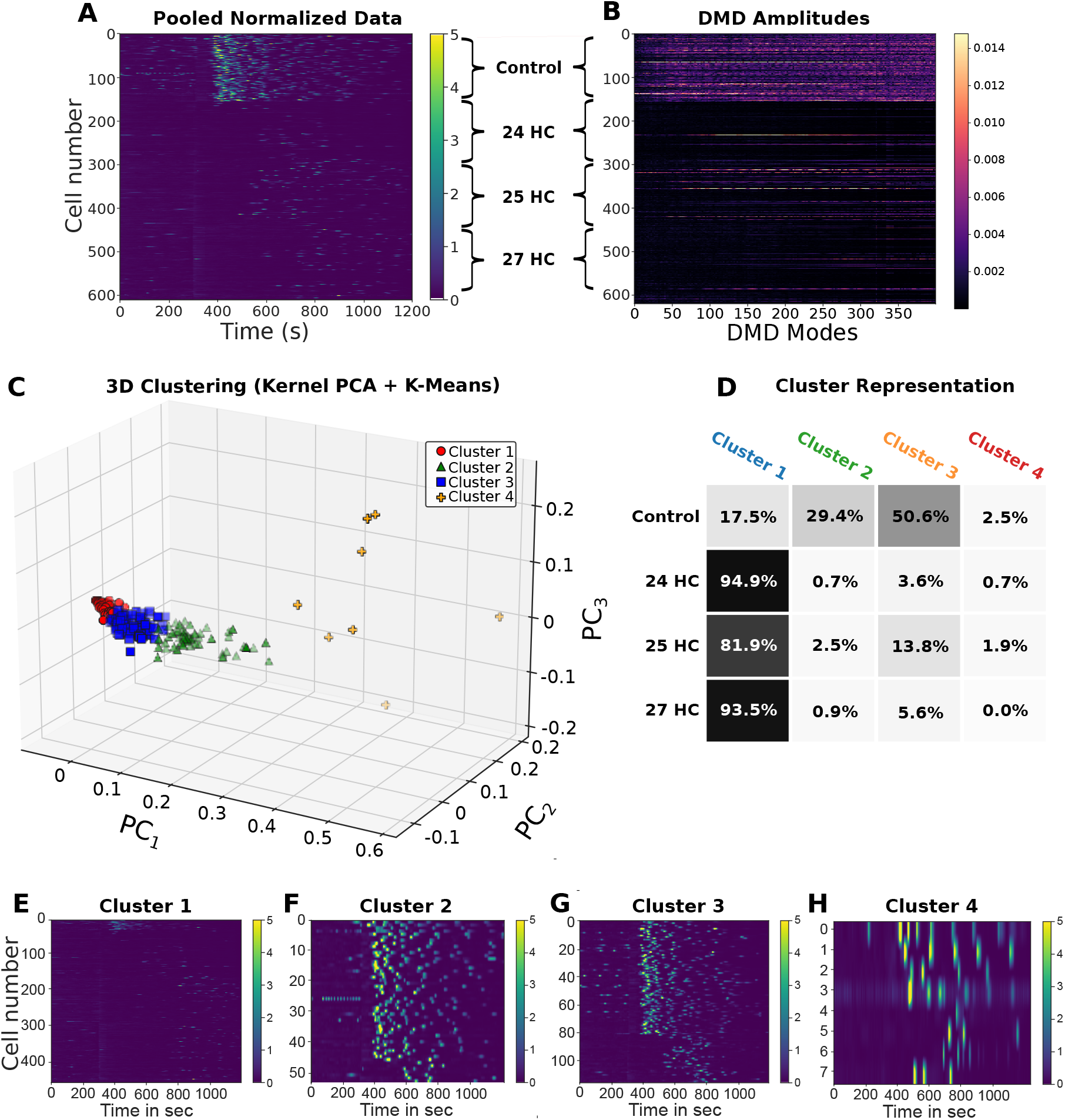
DMD classification of astrocytic Ca^2+^ dynamics after oxysterols treatment and cholesterol loading. (A) Pooled and normalized Ca^2+^ activity traces from all recorded astrocytes across conditions (Control, 24-HC, 25-HC, 27-HC). Each row represents a single cell.(B) Time-delay Dynamic Mode Decomposition amplitudes for all cells, showing the contribution of each DMD mode to individual cellular activity. (C) Three-dimensional embedding of the DMD amplitude features using kernel PCA followed by *k*-means clustering. Four distinct dynamical clusters emerge, separating cells based on shared temporal motifs in their Ca^2+^ activity. Silhouette score = 0.718. (D) Cluster composition across experimental conditions. Oxysterol-treated astrocytes are predominantly assigned to Cluster 1, which is characterized by weak or quiescent Ca^2+^ activity, whereas clusters exhibiting oscillatory or spike-like behavior (Clusters 2 and 3) are enriched in control cells. (E-H) Ca^2+^ activity heatmaps for cells belonging to each cluster. Cluster 1 (E) displays recurrent, temporally coherent activity. Cluster 2 (F) shows sparse or unstructured signals with minimal coordination. Cluster 3 (G) exhibits pronounced bursting patterns, while Cluster 4 (H) contains a small subset of cells with irregular or atypical Ca^2+^ dynamics. Together, these results demonstrate that DMD-TDE combined with unsupervised clustering reveals distinct dynamical phenotypes of astrocytic Ca^2+^ signaling and highlights condition-dependent shifts in astrocyte activity patterns.

Time-delay DMD revealed clear differences in the temporal structure of Ca^2+^ activity across conditions (Fig. 6B). Kernel PCA followed by *k*-means clustering separated the singlecell traces into four distinct activity classes (Fig. 6C). However, the distribution of cells across these clusters differed substantially from that observed in the cholesterol-only experiments (Fig. 6D). Cells pretreated with oxysterols were predominantly assigned to cluster 1, characterized by weak or quiescent Ca^2+^ activity, whereas clusters 2-4, which exhibited oscillatory or spike-like behavior, were enriched in control cells. Among the oxysterol-treated conditions, only a small fraction of 25-HC-treated cells appeared in cluster 3.

Representative traces from each cluster further illustrate these differences in Ca^2+^ dynamics (Fig. 6E-H). Notably, none of the oxysterols reproduced the Ca^2+^ activity pattern induced by cholesterol loading alone. Instead, pretreatment consistently shifted astrocytes toward dampened or inactive Ca^2+^ signaling states.

Consistent with these observations, peak-based analysis showed a reduction in Ca^2+^ spike frequency following oxysterol pretreatment. While such metrics capture overall changes in activity, they provide only a coarse description of the underlying single-cell dynamics. In particular, peak counts and Δ*F/F*_0_ values do not distinguish between sparse high-amplitude events and dense low-amplitude oscillations. By contrast, DMD-TDE resolves these differences by capturing the full temporal structure of the Ca^2+^ signals, allowing distinct activity patterns to be identified across the cell population.

## Discussion

In this study, we combined large-scale single-cell Ca^2+^ imaging with DMD-based time series analysis to characterize how changes in cholesterol influence astrocytic Ca^2+^ signaling. By using both simulated data and experimental recordings from human astrocytes, we show that this approach can dissect heterogeneous Ca^2+^ signals into distinct clusters and relate these clusters to defined changes in sterol loading. Our results indicate that the cholesterol levels in the plasma membrane are crucial for the downstream Ca^2+^ signaling.

The simulations based on a well-established ODE model of astrocyte Ca^2+^ signaling served as a controlled test case for our analysis. In the simulations, variation of SERCA activity was sufficient to generate a broad range of oscillatory Ca^2+^ behaviors that resemble those observed experimentally. Although SERCA activity was not directly manipulated in the present experiments, previous work has shown that pharmacological inhibition of SERCA using thapsigargin abolishes cholesterol-induced Ca^2+^ oscillations in astrocytes (56).

Applying this framework to human astrocytes revealed clear effects of cholesterol loading and cholesterol depletion on Ca^2+^ signaling. Under control conditions, many cells showed spontaneous Ca^2+^ oscillations with occasional spikes, in line with previous reports of basal astrocytic activity (57). Depleting cholesterol with cyclodextrin almost completely removed this activity, whereas loading cholesterol increased the height and frequency of the Ca^2+^ oscillations, especially at 100 and 250 µM cholesterol–cyclodextrin. DMD-TDE identified modes that changed in a concentrationdependent manner, and clustering of DMD modes separated cells into three groups with distinct Ca^2+^ responses. Cells exposed to high cholesterol were mainly found in clusters with strong and frequent spiking, whereas control and cholesterol-depleted cells were enriched in a cluster with weak or absent Ca^2+^ activity. Thus, the levels of cholesterol in the membrane seems to correlate to the astrocytic Ca^2+^ activity.

We next examined whether cholesterol loading affects the coordination of Ca^2+^ activity across astrocytes. Both temporal correlation analysis and the Kuramoto order parameter revealed an increase in synchronized Ca^2+^ activity shortly after cholesterol addition, but only within specific clusters, which were identified by DMD-TDE. By comparison, cells that were common under control or cholesterol-depleted conditions displayed largely stable and weakly coordinated activity. These findings suggest that cholesterol does not only influence Ca^2+^ amplitude and frequency at the single-cell level, but may also facilitate coordinated activity among groups of astrocytes, which could be important for networklevel signaling and communication with neurons.

A central result of this work is that oxysterols strongly changed the way astrocytes respond to acute cholesterol loading. Pretreatment with 24-HC, 25-HC, or 27-HC for 24 hours nearly abolished the large, coordinated Ca^2+^ response normally seen after adding cholesterol-cyclodextrin. DMDTDE analysis revealed four clusters, but in contrast to the cholesterol-only experiments, oxysterol-treated cells accumulated mainly in a cluster with low or quiescent Ca^2+^ activity. Clusters with pronounced oscillatory or spike-like behavior were dominated by control cells. These observations demonstrate that oxysterols reduce the Ca^2+^ response to acute cholesterol loading. One possible explanation is that oxysterol pretreatment changes the pool of accessible cholesterol at the plasma membrane before cholesterol-cyclodextrin is added. In other cell-lines, 25-hydroxycholesterol has been shown to deplete accessible plasma-membrane cholesterol through a mechanism involving ACAT-dependent esterification and suppression of SREBP signaling (58). In astrocytes, 24(S)-hydroxycholesterol can also activate LXR-dependent pathways that promote cholesterol efflux, including apoE-mediated export (59), which could similarly reduce the membrane cholesterol pool available for acute responses. In this context, our finding that oxysterol pretreatment largely abolishes the cholesterol-induced Ca^2+^ response extends earlier observations that cholesterol loading can trigger robust astrocytic Ca^2+^ oscillations (15), and suggests that the ability of astrocytes to respond depends on their sterol state prior to cholesterol delivery.

Our approach also provides a general pipeline for analyzing heterogeneous Ca^2+^ signals at scale. Traditional analyses based on peak counting, average ΔF/F, or manual inspection can easily miss rare or intermediate behaviors, especially when hundreds of cells are recorded simultaneously. In contrast, DMD-TDE uses the full time course and captures both amplitude and temporal structure of the Ca^2+^ signals. Kernel PCA and k-means clustering then group cells according to their temporal modes rather than arbitrary thresholds. This allowed us to identify clusters with high-frequency spiking, broad isolated peaks, damped oscillations, or almost flat traces, and to relate these clusters to specific experimental conditions. The same strategy can in principle be applied to other cell types, stimuli, or imaging modalities, making it a versatile tool for studying complex Ca^2+^ dynamics. We also tested an alternative trajectory clustering method, which is based on calculating principal angles on the Grassmann manifold (See Fig. S1). This analysis also provided very good clustering outputs, but was limited to the synthetic data. Still this clustering approach could be a useful alternative in future studies.

Although experiments were performed under a limited number of conditions, each dataset contains hundreds of singlecell Ca^2+^ trajectories, enabling robust unsupervised classification of dynamical states based on temporal structure rather than population averages. Accordingly, the emphasis of this study is methodological, focusing on the ability of DMD-TDE to resolve heterogeneous signaling behaviors, rather than on population-level statistical inference across multiple biological replicates.

## Conclusion

In this study, we established a high-throughput experimental and computational framework for quantifying the heterogeneity of astrocytic Ca^2+^ dynamics and for determining how changes in sterol composition shape these signaling states. By combining widefield Ca^2+^ imaging with deeplearning–based single-cell segmentation and DMD-TDE, we were able to resolve distinct temporal modes of Ca^2+^ activity across hundreds of individual astrocytes simultaneously. This approach successfully classified both simulated and experimental data, capturing oscillatory, transient, and quiescent behaviors that are otherwise difficult to distinguish with conventional analysis methods.

Using this framework, we found that acute cholesterol loading produces strong, coordinated Ca^2+^ elevations and distinct oscillatory signatures, whereas cholesterol depletion markedly suppresses Ca^2+^ activity. Pretreatment with hydroxycholesterols, however, profoundly altered these responses, shifting the population toward dampened and largely quiescent signaling states. These findings demonstrate that even closely related sterol derivatives exert highly specific effects on astrocytic Ca^2+^ dynamics, highlighting the sensitivity of Ca^2+^ signaling to membrane composition and lipid homeostasis.

By resolving Ca^2+^ heterogeneity across large astrocyte populations, this framework offers new opportunities to uncover how sterol metabolism, membrane organization, and intracellular Ca^2+^ handling intersect to regulate astrocyte physiology and potentially contribute to disease-associated signaling states.

## Acknowledgements

The work presented here is supported by the Carlsberg Foundation (grant CF23-1086) and the Lundbeck Foundation (grant no. R366-453 2021-226). We would further like to acknowledge the imaging facility DaMBIC for providing microscopy infrastructure (Novo Nordisk Foundation, grant no. NNF18SA0032928).

## Author Contributions

ML has carried out all experiments and analyzed the data with help from LL and DW. ML has also written large parts of the manuscript with input and feedback from all authors. DWhas designed the study, developed the model and raised funding for the project. RJ and RZ have designed and implemented the Grassmann-based clustering and given feedback on the background theory. All authors have read and approved the manuscript.

**ACKNOWLEDGEMENTS**

