## Supplementary for "Single-cell analysis of sterol-induced Ca^2+^ signaling in human astrocytes by dynamic mode decomposition"

### Supplementary Note 1:

**A. Clustering of DMD bases on the Grassmann manifold.** In this section, we describe an alternative clustering strategy based on comparing delay-DMD bases on the Grassmann manifold. While this approach performed well on synthetic data generated from the ODE model used throughout this study (see below), we were unable to obtain reliable results when applying it to individual experimental $Ca^{2+}$ traces. In practice, DMD-TDE applied to single experimental time series frequently yielded unstable or poor reconstructions due to the combined effects of high noise levels, high effective dimensionality, and sparse temporal sampling of the experimental data. Because the method relies on stable low-dimensional subspace representations for each time series, these properties limit its applicability to the simulated datasets considered here.

Given a collection of time series $\{x_i(t) : i \leq M\}$, DMD-TDE is applied individually to snapshot data

$$[x_i(t_0), \ldots, x_i(t_m)].$$

For fixed delay dimension $d$ and truncation rank $r$, this yields a set of DMD bases $\{\Phi_1, \ldots, \Phi_M\}$, where each basis $\Phi_i \in \mathbb{R}^{d\times r}$ spans an $r$-dimensional subspace of $\mathbb{R}^d$ and thus represents a point on the Grassmann manifold $\mathrm{Gr}(d,r)$ (60, 61). We apply Grassmann spectral clustering (62) to assess whether differences in the underlying dynamics can be detected at the level of these subspaces.

To compare two DMD bases $\Phi_i$ and $\Phi_j$ ($i \neq j$), thin QR decompositions are computed to obtain orthonormal bases $Q_i$ and $Q_j$. Their distance is defined using the principal angles between the corresponding subspaces,

$$d(Q_i, Q_j)^2 = \sum_{k=1}^{r} \arccos(\sigma_k)^2, \qquad Q_i^T Q_j = U \operatorname{diag}(\sigma_1, \ldots, \sigma_r) V^T. \tag{10}$$

Based on this distance, the affinity matrix $A$ for spectral clustering is defined as

$$[A]_{ij} = \exp\left(-\frac{d(Q_i, Q_j)^2}{\sigma^2}\right), \tag{11}$$

with kernel parameter $\sigma$ and zero diagonal entries. Clustering into $K$ groups is performed using Algorithm 2 of (62).

We validated this approach using synthetic $Ca^{2+}$ time series generated from the astrocyte ODE model described in (45). To introduce controlled variability in the dynamics, the SERCA-associated parameter $v_{m2}$ was varied around four mean values $\mu_i = 2, 11, 14, 17$ by adding Gaussian noise drawn from a standard normal distribution, with ten independent realizations generated for each mean. The ODE system was integrated using `ode15s` with initial condition $x_0 = (0,0,0)^T$, while all remaining parameters were fixed as in (45). Each resulting trajectory $X_k(t)$ ($k = 1, \ldots, 40$) was sampled at $t = 0, 1, \ldots, 400$, and a DMD-TDE basis was computed using delay dimension $d = 200$ and truncation rank $r = 40$.

Grassmann spectral clustering was applied with $K = 3$ clusters and kernel parameter $\sigma^2 = 70$. The resulting clustering (Fig. S1) separates the time series into three groups corresponding closely to the parameter centers $\mu_i$, demonstrating that this approach can distinguish parameter-dependent dynamical regimes in controlled synthetic data.

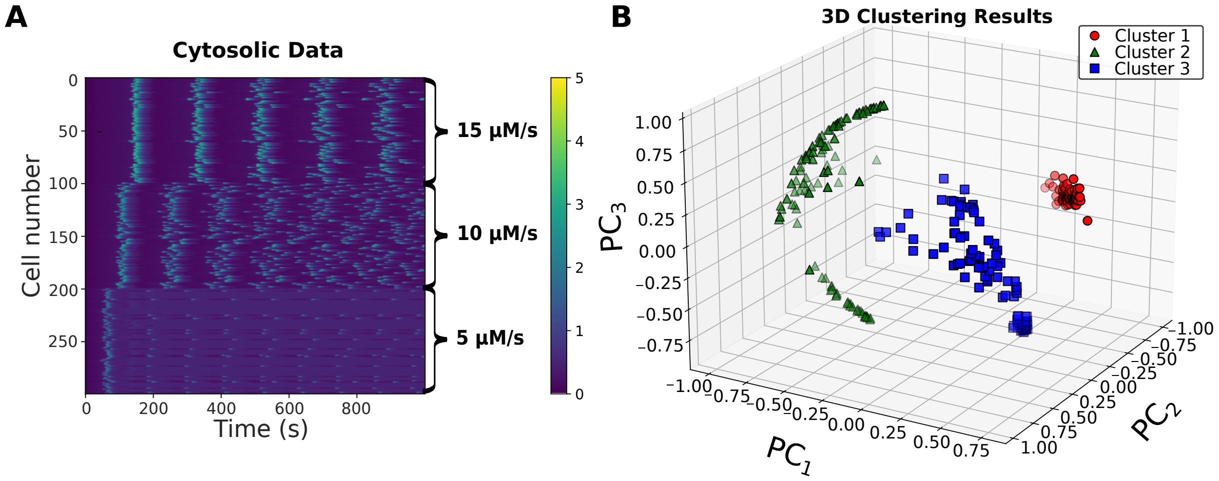


**Fig. S1.** Clustering of DMD-TDE bases on the Grassmann manifold. (A) Synthetic $Ca^{2+}$ time series generated from the astrocyte ODE model for representative values of the SERCA-associated parameter $v_{m2}$. (B) Distribution of cluster assignments obtained by Grassmann spectral clustering of DMD-TDE bases. For each value of $v_{m2}$, the number of trajectories assigned to each cluster is shown, demonstrating separation of dynamical regimes across parameter values.

***Temperature modulates spontaneous Ca$^{2+}$ activity in astrocytes.*** Temperature influences biochemical reaction rates, membrane fluidity, and ion channel kinetics, all of which are central determinants of intracellular Ca$^{2+}$ signaling. We therefore asked whether imaging temperature alone is sufficient to alter spontaneous astrocytic Ca$^{2+}$ dynamics under otherwise identical experimental conditions.

Astrocytes imaged at 37 °C exhibited increased spontaneous Ca$^{2+}$ activity compared to cells imaged at 20 °C (Fig. S2). Quantitative analysis revealed an increase in the number of Ca$^{2+}$ spikes per cell and a larger fraction of active cells at 37 °C (Fig. S2C-D). In addition, Ca$^{2+}$ transients at 37 °C were shorter in duration and of lower peak amplitude compared to those observed at 20 °C (Fig. S2E-F). Together, these results indicate that temperature reshapes multiple features of spontaneous astrocytic Ca$^{2+}$ signaling.

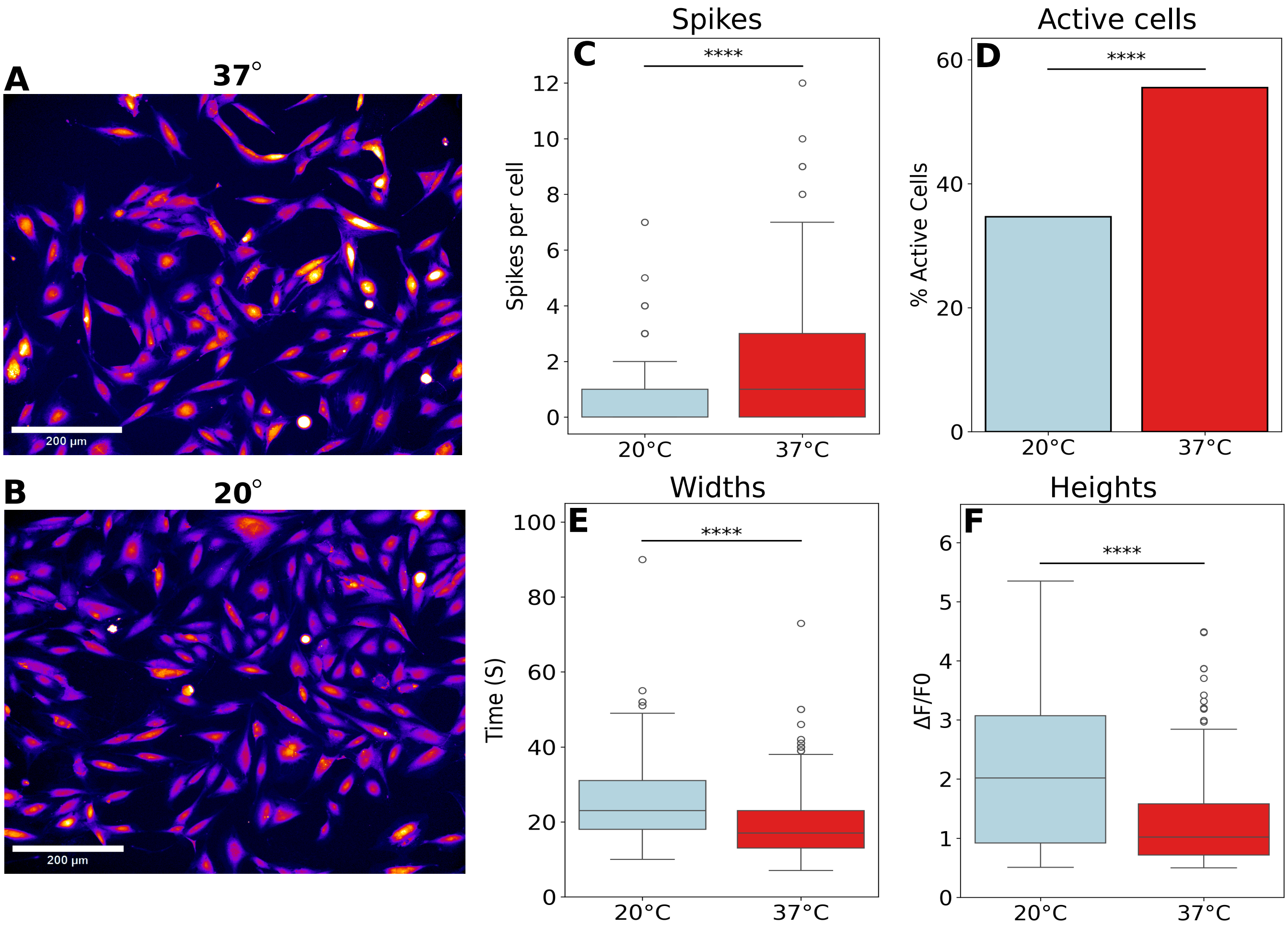


**Fig. S2.** Temperature-dependent modulation of spontaneous astrocytic Ca$^{2+}$ activity. (A,B) Representative fluorescence images of astrocytes loaded with Cal-520 and imaged at 37 °C (A) or 20 °C (B). Scale bars: 200 $\mu$m. (C) Number of Ca$^{2+}$ spikes per cell. (D) Percentage of active cells exhibiting at least one Ca$^{2+}$ transient during the recording period. (E) Spike width (full width) of Ca$^{2+}$ transients. (F) Peak amplitude of Ca$^{2+}$ transients expressed as $\Delta F/F_0$. Statistical significance was assessed using [test], with **** indicating $p < 0.0001$; sample sizes are indicated in the panel labels.

**Cholesterol availability modulates astrocytic $Ca^{2+}$ activity.** We compared $Ca^{2+}$ activity in control cells, cells treated with MCD alone, and cells exposed to increasing concentrations of cholesterol-MCD complexes. This design allows direct comparison of cholesterol depletion and loading under identical imaging conditions.

Cholesterol depletion by MCD alone significantly reduced spontaneous $Ca^{2+}$ activity relative to control cells, as reflected by fewer $Ca^{2+}$ spikes per cell (Fig. S3A). In contrast, cholesterol loading produced a concentration-dependent increase in $Ca^{2+}$ spike frequency, with the highest cholesterol condition (250 $\mu$M) exhibiting the greatest number of spikes. Cholesterol loading also increased $Ca^{2+}$ spike amplitudes (Fig. S3B), whereas cholesterol depletion did not significantly alter spike heights relative to control. Spike durations were largely unchanged across conditions, with no significant differences detected relative to control (Fig. S3C).

Analysis of the cumulative distribution of first-spike times revealed that cholesterol depletion delayed the onset of $Ca^{2+}$ activity, whereas cholesterol loading accelerated cellular responsiveness in a dose-dependent manner (Fig. S3D). Together, these results demonstrate that cholesterol availability modulates both the magnitude and timing of astrocytic $Ca^{2+}$ signaling.

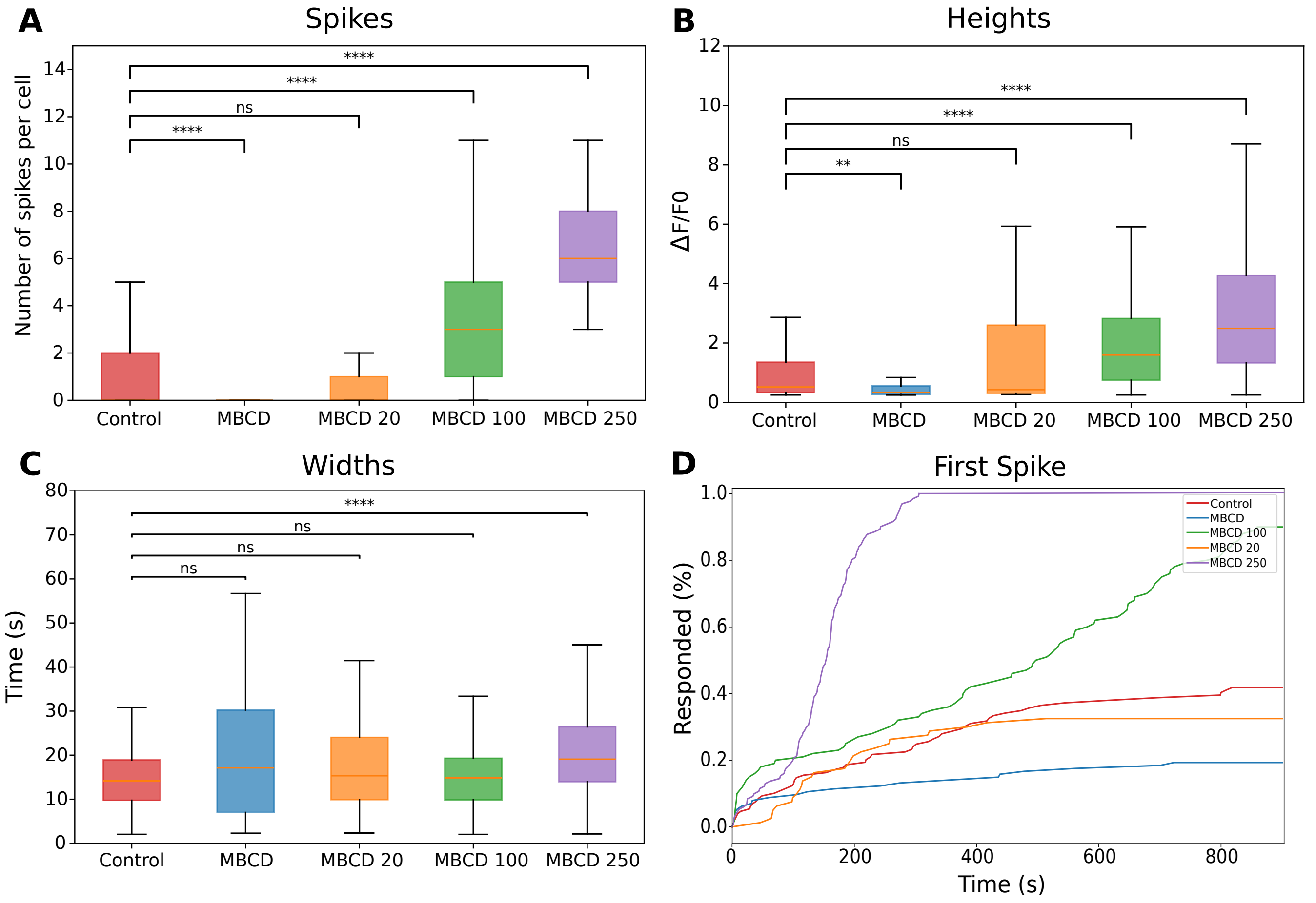


**Fig. S3.** Cholesterol loading and depletion modulate spontaneous $Ca^{2+}$ activity in human astrocytes. Astrocytes were imaged using Cal-520 during live-cell $Ca^{2+}$ imaging and treated with control solution, MCD alone, or cholesterol-MCD complexes (20-250 $\mu$M cholesterol) after 300 s of baseline recording. $Ca^{2+}$ activity was extracted on a per-cell basis and normalized as $\Delta F/F_0$. A total of 129 control cells (159 $Ca^{2+}$ peaks), 114 MCD-treated cells (34 peaks), 80 cholesterol-MCD 20 $\mu$M cells (54 peaks), 100 cholesterol-MCD 100 $\mu$M cells (381 peaks), and 131 cholesterol-MCD 250 $\mu$M cells (888 peaks) were analyzed. (A) Number of $Ca^{2+}$ spikes per cell. MCD treatment reduced spike counts relative to control, whereas cholesterol loading produced a concentration-dependent increase. (B) $Ca^{2+}$ spike amplitudes ($\Delta F/F_0$). Spike amplitudes were reduced following MCD treatment and increased at higher cholesterol concentrations. (C) $Ca^{2+}$ spike widths. Spike durations were largely unchanged across conditions. (D) Cumulative distribution of first-spike times showing the fraction of responding cells over time. MCD treatment delayed and reduced $Ca^{2+}$ activation, whereas cholesterol loading accelerated cellular responsiveness in a dose-dependent manner. Statistical significance is indicated as not significant (ns), * $p < 0.05$, ** $p < 0.01$, *** $p < 0.001$, **** $p < 0.0001$.

***Kinetics of sterol delivery by MCD.*** Because cholesterol-induced $Ca^{2+}$ responses occur rapidly after addition of cholesterol-MCD complexes, we verified the kinetics of sterol delivery under the imaging conditions used in this study. To directly visualize sterol uptake, we used DHE, a naturally fluorescent cholesterol analog that closely mimics cholesterol membrane partitioning and intracellular trafficking.

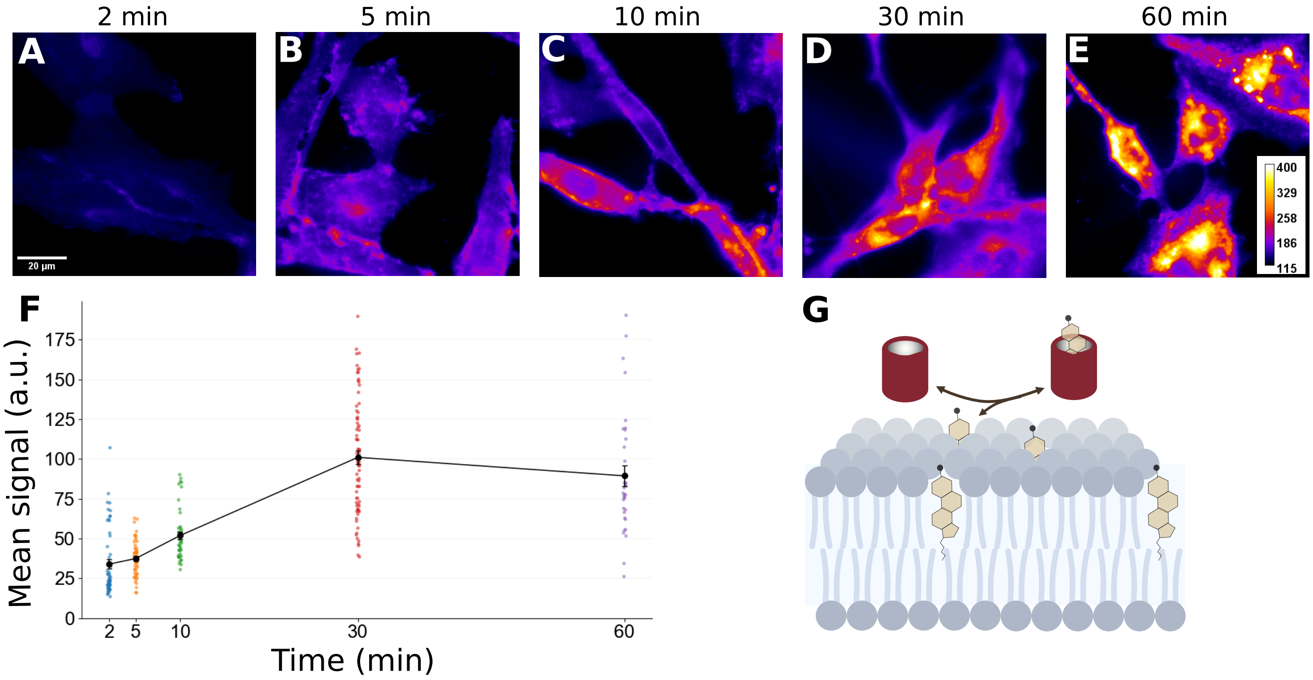


**Fig. S4.** Kinetics of sterol delivery by DHE-MCD complexes. Astrocytes were exposed to MCD loaded with 250 $\mu$M DHE for 2, 5, 10, 30, or 60 min to monitor sterol delivery dynamics. (A-E) Representative fluorescence images showing progressive accumulation of DHE over time. (F) Quantification of mean cellular DHE fluorescence per cell across time points, demonstrating rapid uptake between 2 and 10 min, a peak at 30 min, and sustained signal at 60 min. Each point represents a single cell. (G) Schematic illustration of sterol delivery by DHE-MCD complexes, in which MCD shuttles sterol molecules to the plasma membrane for incorporation into the lipid bilayer.

**DMD-based clustering of cholesterol-induced $Ca^{2+}$ responses.** Single-cell $Ca^{2+}$ traces obtained after stimulation with cholesterol-MCD complexes were analyzed using DMD-TDE followed by kernel PCA and k-means clustering. This analysis separated astrocytes into three response clusters (Clusters 1-3) based on the temporal structure of their $Ca^{2+}$ activity.
Cluster 1 was characterized by weak or absent $Ca^{2+}$ activity. Cluster 2 cells exhibited sporadic, high-amplitude $Ca^{2+}$ spikes, whereas Cluster 3 cells displayed more frequent, lower-amplitude oscillatory activity. Quantification of spike number, amplitude, width, and first-spike timing revealed significant differences between clusters, indicating distinct $Ca^{2+}$ signaling phenotypes in response to cholesterol loading.

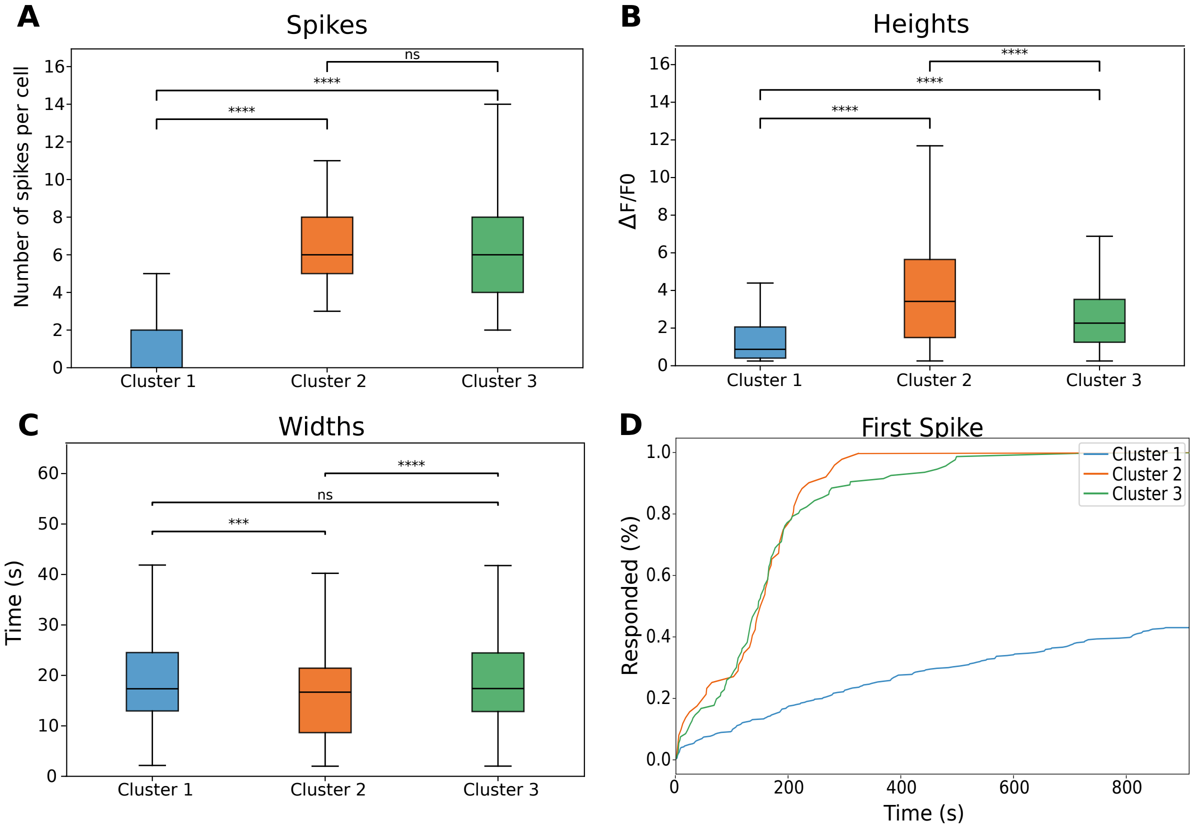


**Fig. S5.** DMD-based clustering of cholesterol-induced $Ca^{2+}$ responses in human astrocytes. Single-cell $Ca^{2+}$ traces obtained after stimulation with cholesterol-MCD complexes were classified into three clusters using DMD-TDE followed by kernel PCA and k-means clustering. (A) Number of $Ca^{2+}$ spikes per cell across clusters. Cluster 1 shows minimal spiking activity, whereas Clusters 2 and 3 exhibit increased spike counts. (B) Spike amplitudes ($\Delta F/F_0$) for each cluster. (C) Spike widths across clusters. (D) Cumulative distribution of first-spike times showing the fraction of responding cells over time. Statistical significance is indicated as not significant (ns), * $p < 0.05$, ** $p < 0.01$, *** $p < 0.001$, **** $p < 0.0001$.

**Oxysterol pretreatment reduces cholesterol-induced $Ca^{2+}$ activity.** To determine whether oxysterols modulate astrocytic $Ca^{2+}$ responses to acute cholesterol loading, astrocytes were pretreated for 24 h with 24-HC, 25-HC, or 27-HC prior to stimulation with cholesterol-MCD complexes during live-cell $Ca^{2+}$ imaging. Oxysterol pretreatment reduced the magnitude and frequency of cholesterol-induced $Ca^{2+}$ activity, indicating that prior sterol exposure dampens astrocytic responsiveness to acute cholesterol delivery.

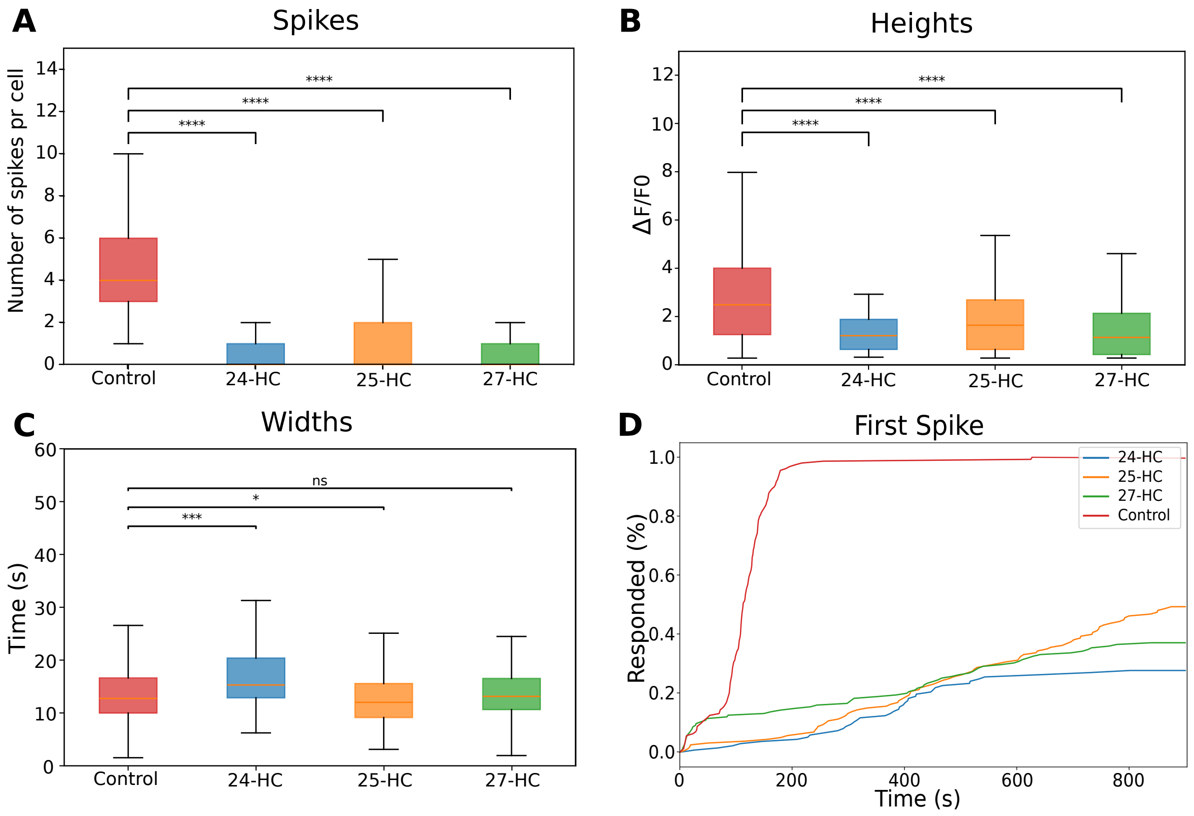


**Fig. S6.** Oxysterol pretreatment attenuates cholesterol-induced $Ca^{2+}$ activity in human astrocytes. Astrocytes were pretreated for 24 h with 24-HC, 25-HC, or 27-HC and imaged using Cal-520 during live-cell $Ca^{2+}$ imaging. Cholesterol-MCD complexes were added after 300 s of baseline recording. $Ca^{2+}$ activity was extracted on a per-cell basis and normalized as $\Delta F/F_0$. A total of 160 control cells (748 $Ca^{2+}$ peaks), 137 24-HC-treated cells (52 peaks), 160 25-HC-treated cells (159 peaks), and 175 27-HC-treated cells (132 peaks) were analyzed. (A) Number of $Ca^{2+}$ spikes per cell. Oxysterol pretreatment reduced spike counts compared to control. (B) $Ca^{2+}$ spike amplitudes ($\Delta F/F_0$), showing attenuated responses in oxysterol-treated groups. (C) $Ca^{2+}$ spike widths, indicating modest changes in calcium signal kinetics following oxysterol pretreatment. (D) Cumulative distribution of first-spike times showing delayed and reduced $Ca^{2+}$ activation in oxysterol-treated astrocytes compared to control. Statistical significance is indicated as not significant (ns), * $p < 0.05$, ** $p < 0.01$, *** $p < 0.001$, **** $p < 0.0001$.
